# Optimized production and fluorescent labelling of SARS-CoV-2 Virus-Like-Particles to study virus assembly and entry

**DOI:** 10.1101/2022.03.23.485575

**Authors:** Manon Gourdelier, Jitendriya Swain, Coline Arone, Anita Mouttou, David Bracquemond, Peggy Merida, Saveez Saffarian, Sébastien Lyonnais, Cyril Favard, Delphine Muriaux

**Affiliations:** Institut de Recherche en Infectiologie de Montpellier (IRIM), Université de Montpellier, CNRS UMR9004, Montpellier, France; Department of Physics and Astronomy, Center for Cell and Genome Sciences, University of Utah, Salt Lake City, Utah, United States; CEMIPAI, Université de Montpellier, CNRS UAR3725, Montpellier, France

**Keywords:** SARS-CoV-2, Virus-Like-Particles, Super resolution microscopy, Virus assembly, Virus entry

## Abstract

SARS-CoV-2 is an RNA enveloped virus responsible for the COVID-19 pandemia that conducted in 6 million deaths worldwide so far. SARS-CoV-2 particles are mainly composed of the 4 main structural proteins M, N, E and S to form 100nm diameter viral particles. Based on productive assays, we propose an optimal transfected plasmid ratio mimicking the virus RNA ratio allowing SARS-CoV-2 Virus-Like Particle (VLPs) formation composed of the viral structural proteins M, N, E and S. Furthermore, monochrome, dual-color fluorescent or photoconvertible VLPs were produced. Thanks to live fluorescence and super-resolution microscopy, we quantified VLPs size and concentration. It shows a diameter of 110 and 140 nm respectively for MNE-VLPs and MNES-VLPs with a minimum concentration of 10e12 VLP/ml. SARS-CoV-2 VLPs could tolerate the integration of fluorescent N and M tagged proteins without impairing particle assembly. In this condition, we were able to establish incorporation of the mature Spike in fluorescent VLPs. The Spike functionality was then shown by monitoring fluorescent MNES-VLPs docking and endocytosis in human pulmonary cells expressing the receptor hACE2. This work provides new insights on the use of non-fluorescent and fluorescent VLPs to study and visualize the SARS-CoV-2 viral life cycle in a safe environment (BSL-2 instead of BSL-3). Moreover, optimized SARS-CoV-2 VLP production can be further adapted to vaccine design strategies.

## INTRODUCTION

The Severe Acute Respiratory Syndrome Coronavirus-2 (SARS-CoV-2) is an emerging virus belonging to the *coronaviridae* family and the causative agent of the Coronavirus Disease 2019 (COVID-19). This respiratory virus emerged in China during the month of December 2019 and spread all around the world, leading to the ongoing pandemic declared in March 2020 by the World Health Organization (WHO). Pathogenicity of coronaviruses was previously observed during SARS-CoV and MERS outbreaks, which respectively occurred in 2002 and 2012. Concerning the current pandemic, COVID-19 has affected up to 460 million cases and caused 6 million deaths (WHO). It’s demonstrating the importance of gathering knowledge on SARS-CoV-2 and developing vaccine design strategies. Among the possible strategies, Virus-Like Particles (VLPs) are high potential as they can be efficient and cheap vaccine candidates [1]. VLPs are multi-protein-lipid structures that mimic the organization and conformation of authentic native viruses without containing the viral genome.

To date, non-infectious VLPs have already been optimized using HIV-1 or Influenza structural proteins, allowing to study the entry and the assembly of these viruses in BSL-2 facilities. While the minimal requirement for HIV-1 VLPs using only one viral protein, i.e. Gag [2], Kerviel and collaborators showed that the Influenza minimal system requires at least four proteins for VLPs production [3]. SARS-CoV-2 is an enveloped virus containing one RNA molecule and 29 proteins, including four structural proteins: membrane protein M, nucleoprotein N, envelope protein E and spike protein S [4]. Thus, determining the optimal combination (and proportion) of these structural viral proteins remains to be tested to obtain a minimal system for SARS-CoV-2 VLPs production.

Recently, research teams establish several minimal systems for SARS-CoV-2 VLPs production [5] [6] [7] [8] [9] [10] [11] exhibiting differences and contradictory results regarding the required structural viral proteins. Hence, different combinations regarding M, N, E and S were reported: the minimal system for VLPs production could require both M and E [5] or M, E and S [7] or M and E with N which plays a crucial role for VLPs formation [10] as previously reported for SARS-CoV [12]. A combination of M with N or S was also suggested [6]. Here, based on years of experiences in the production of HIV-1 Gag VLPs and their characterization using advanced microscopies [11] [12] [15] [16], our work aims to revisit the minimal system for SARS-CoV-2 VLPs production. To optimize VLPs production from DNA plasmid ratio of M, N, E or S, we based our approach on viral RNA ratios obtained from transcriptomic analysis of SARS-CoV-2 RNA genome in infected cells [17] [18]. We also optimized the production of fluorescent VLPs (mono- or bi-color), made with tagged structural SARS-CoV-2 proteins, to create a valuable tool to image and study SARS-CoV-2 assembly and entry. We characterize these VLPs thanks to advanced microscopy methods such as Total Internal Reflection Fluorescence Microscopy (TIRF-M), Fluctuation Correlation Spectroscopy (FCS), correlative fluorescence-Atomic Force Microscopy (AFM) and Photo-Activable Localization Microscopy (PALM). These techniques allow us to determine the size, numbers and morphology of VLPs that we could compare to wild-type SARS-CoV-2 particles [18] [20]. Finally, these fluorescent VLPs, made with tagged structural proteins (M or N) allowed us to evaluate the incorporation of the Spike and to follow viral particle entry into host pulmonary cells using quantitative confocal microscopy.

## MATERIALS AND METHODS

### Cells and culture conditions

The HEK293T human embryonic kidney cell line were obtained from ECACC (Sigma-Aldrich, Germany) and maintained in cultured in Dulbecco’s modified essential medium (DMEM, Gibco) supplemented with 10% heat inactivated fetal calf serum (FCS, Thermo Fisher, USA), 50 U/mL of penicillin (Ozyme, France), 50 μg/mL of streptomycin (Ozyme, France), 1mM sodium-pyruvate (Ozyme, France) and 25mM of HEPES (Ozyme, France), at 37 °C with 5% CO_2_.

Pulmonary A549 adenocarcinoma human alveolar basal epithelial cell lines [13] were obtained from ECACC (#86012804, Sigma-Aldrich, Germany) and cultured in Roswell Park Memorial Institute medium (RPMI) from Gibco supplemented with 10% heat inactivated fetal calf serum (FCS, Thermo Fisher, USA), 50 U/mL of penicillin (Ozyme, France), 50 μg/mL of streptomycin (Ozyme, France), 1mM sodium-pyruvate (Ozyme, France) and 25mM HEPES (Ozyme, France), at 37°C with 5% CO_2_. The human pulmonary alveolar A549-hACE2 and A549-hACE2mScarlet stable cell lines were engineered using 2 different lentiviral vectors. For A549-hACE2, A549 cells were transduced with VSVg pseudotyped particles derived from lentiviral vector from Flash Therapeutics (Toulouse, France) to express the human receptor ACE2. Cells were then sorted by flow cytometry (BD FACSAria™ III Cell Sorter) in order to obtain an A549-hACE2 population stably expressing hACE2. To detect hACE2, cells were washed twice in PBS and incubated 1 hour at 4°C with primary goat antibody anti-human ACE2 protein (R&D AF933) (1:100). After washing in PBS, cells were stained 45min at 4°C with secondary antibody AF488-labeleddonkey-anti-goat (Lifetechnologies A11055) (1:3000) for 45min. h-ACE2 expression was monitored in comparison with isotype control using NovoCyte Flow Cytometer (AGILENT) (Supplemental Figure 1A). For A549-hACE2-mScarlet, A549 were transduced with VSVg pseudotypes particules derived from lentiviral vector expressing mScarlet red fluorescent protein from Olivier Schwartz laboratory at Institut Pasteur (Paris, France) to express the human receptor hACE2mScarlet. Transduction efficiency was assessed by monitoring mScarlet expression (Maximum Excitation 569nm/Maximum Emission 594nm) using flow cytometry in comparison with A549 cells (Supplemental Figure 1B).

### DNA plasmids and cloning

The plasmid expressing Wuhan SARS-CoV-2 M protein, N protein, E protein and S protein were humanized and put under the control of the cytomegalovirus (CMV) promoter driven mammalian expression vectors as described in Swann et al. [7]. The SARS-CoV-2 M protein was tagged at the C-terminus placing the fluorescent protein within the lumen of the SARS-CoV-2 VLPs. To generate a functional tag, the following protein linker was used: Flex (Flexible protein linker): GGGGSGGGGSGGGGSGGGGD to generate both M-Flex-mCherry and M-Flex-GFP. The plasmids were generated by synthesis of Flex-mCherry and Flex-eGFP and inserted into humanized M plasmids described in [7] using EZcloning from Genscript. The SARS-CoV-2 N protein was tagged at the N-terminus. The GFP and mEOS2 tags were amplified by PCR from peGFP and pmEOS2 using *Pfu* Polymerase (M7741, Promega), dNTP mix (10297018, Thermo Scientific) and primers designed to carry NheI and KpnI restriction sites (Sigma Aldrich). PCR amplified inserts were added at the N-terminus of N protein in pcDNA3.1+ plasmid, and purified with the NucleoSpin Gel and PCR Clean-up kit (Macherey-Nagel). The plasmid N and inserts were digested and purified. Ligation between these digested products was performed using T4 DNA ligase (M0202S, NEB). 50μl of competent bacteria were next transformed with 5μl of the ligation reaction product. Plasmids were extracted from various colonies with the NucleoSpin plasmid mini-kit (740588.50, Macherey-Nagel). Validated plasmids were then sequenced by Eurofins.

### DNA transfection

HEK293T cells were seeded into 6-well plate or 10 mL dishes 24h before transfection. Plasmids expressing viral proteins were transfected with CaCl_2_/HBS2X (50mM HEPES pH 7.1, 280mM NaCl, 1.5mM Na_2_HPO_4_) (v/v) into HEK293T cells. A total of 1.4μg of mixed plasmids was applied for the transfection of a well (for 6-well plate) and 0.7μg/well for individual plasmid. Co-transfection of double, triple or quadruple plasmids was conducted with M, N, E and S with plasmid ratio 3:3:3:5 respectively (0.3μg:0.3μg:0.3μg:0.5μg per well to 6-well plate for example) in the first assay as shown in Figure 1B. Optimization of SARS-CoV-2 VLPs was performed by co-transfecting M, N, E and S with a plasmid ratio of 3:12:2:5 respectively corresponding to the molecular viral RNA ratio found in infected cells [17] [18], corresponding, for an example, to a total of 2,2μg/well of 6 well plate (with M:N:E:S corresponding to 0.3μg:1.2μg:0.2μg:0.5μg of transfected plasmid) as shown in Figure 1C. Optimization of fluorescent SARS-CoV-2 VLPs production was performed by co-transfecting M, M(GFP) or M(mCherry), N, E and S with plasmid ratio of 2:2:12:2:5 respectively, and M, N, N(GFP) or N(mEOS2), E and S with plasmid ratio of 3:9:3:2:5 respectively. Then, transfected cells were washed 6h later with phosphate-buffered saline (PBS 1x) and harvested 24h to 48h post-transfection with Exofree medium. Attached cells were washed with PBS, harvested, centrifuged at 1500 rpm for 5 minutes at 4°C and lysed with RIPA lysis buffer (25mM Tris HCl pH 7.6, 150 mM NaCl, 1% NP-40, 1% sodium deoxycholate, 0.1% SDS) (ThermoFisher) supplemented with protease inhibitor cocktail (Sigma). Following centrifugation at 15 000 rpm for 10 minutes at 4°C, cell lysates were collected. The protein concentrations in cell lysates were estimated using Bradford assays. Under each condition, the percentage of cell viability was measured by trypan blue cell counting before cell lysis.

**Figure 1:**
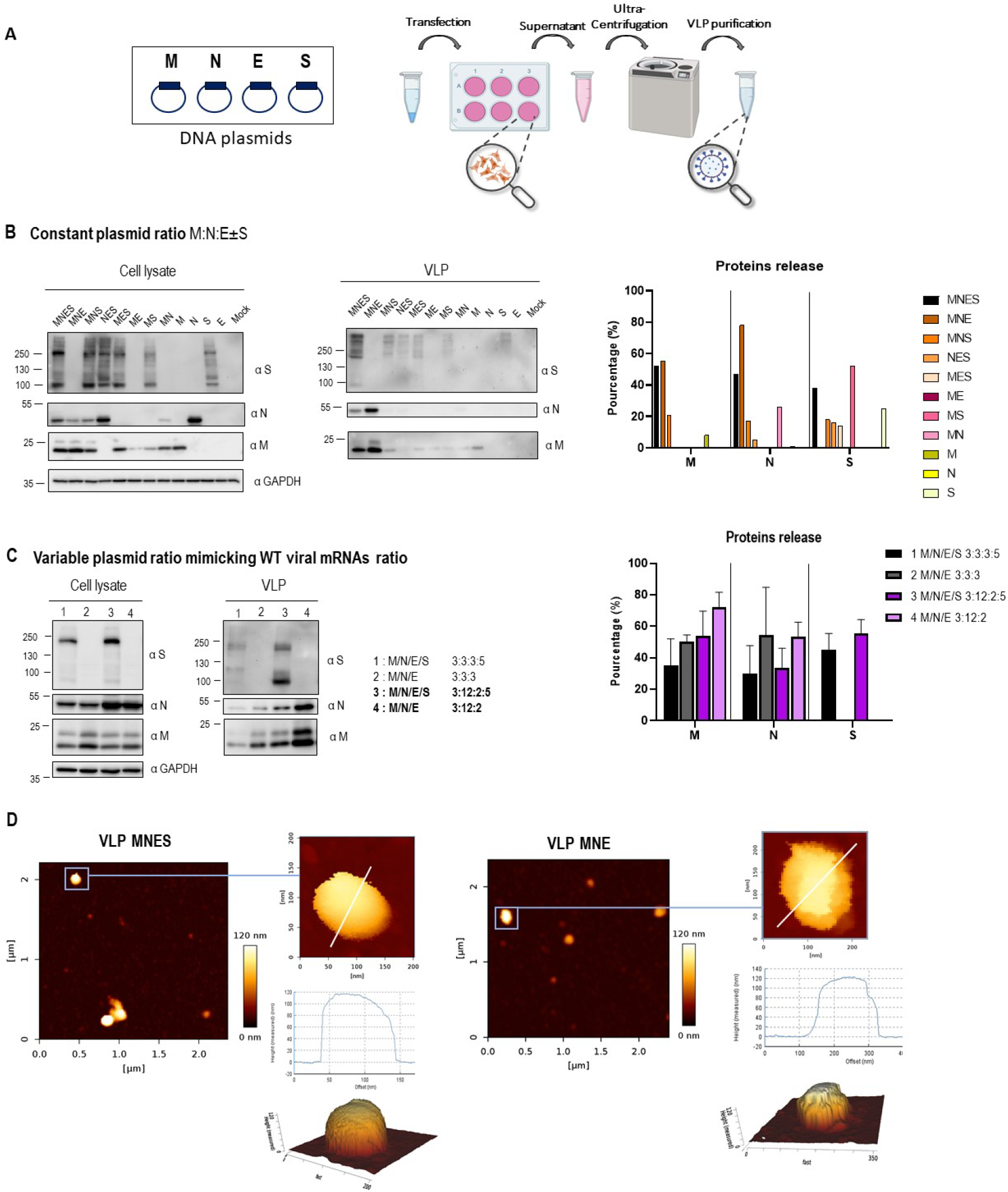
Optimization of SARS-CoV-2 VLPs assembly and particle analysis using immunoblots and atomic force microscopy imaging. (A) Scheme of VLPs production and purification. (B) Western blot of cell lysates and VLPs of HEK293T cells transfected with M, N, E or S as indicated for each lane. The transfected M/N/E/S plasmid ratios are respectively MNES 3:3:3:5; MNE 3:3:3; MNS: 3:3:5, NES 3:3:5, MES 3:3:5, ME 3:3, MS 3:5, M 3, N 3, E 3, S 5. Right panel: percentage of M, N, E and S in released VLPs after quantification of the band intensities on the Western Blot for each condition. (C) Western blot of cell lysates and VLPs of HEK293T cells transfected with the indicated 3:3:3: ± 5 and 3:12:2: ±5 M, N, E ± S transfected plasmid ratios. Right panel: percentage of M, N, E and S in released VLPs after quantification of the western blot band intensities for each condition. N=3 experiments. (D) AFM imaging of SARS-CoV-2 MNES VLPs and MNE VLPs using quantitative QI mode AFM in TNE buffer. For each condition, a topographic image of 2.5 μm x 2.5 μm and a zoom of a particle is shown, with a 3D projection. The color gradient for Z scale is the same for all panels. The topographical profile plots along the white line are shown for each VLP.

### Collected VLPs and purification

VLPs were obtained from culture medium of transfected cells by clarification at 5000 rpm for 5 min at 4°C. Then the VLPs were collected from clarified supernatants after ultracentrifugation, through a 25% sucrose cushion in TNE buffer (10 mM Tris-HCl pH 7.4, 100 mM NaCl, 1 mM EDTA), at 28 000 rpm or 30 000 rpm for 3h at 4°C in Beckman SW41Ti or SW55Ti rotors respectively depending on the volume. VLPs from ultracentrifugation were resuspended overnight at 4°C in TNE buffer. The purified VLPs were stored at 4°C.

### Antibodies

Immunoblotting were performed by using the following antibodies: rabbit anti-M 1:1000 (100-401-A55, Rockland TEBU-BIO);rabbit anti-N 1:3000 (200-401-A50, Rockland TEBU-BIO); mouse anti-S 1:1000 (GTX632604, Genetex); anti-GAPDH antibody coupled to HRP 1:25000 (G9295, Sigma). For immuno-spotting, rabbit anti-Spike neutralizing Antibody 1:100 (40592-R001, Sino Biological), mouse anti-CD81 1:100 (sc7637, 1.3.3.22, Santa Cruz), fluorescent rabbit Alexa555 and mouse Alexa647-conjugated secondary antibodies 1:2000 (Invitrogen) were used in this study.

### Immunoblotting

Proteins from cell lysates (20μg of total proteins) or VLPs (volume to volume) were loaded and separated on 8% and 12% SDS-PAGE gel and transferred onto a polyvinylidene difluoride transfer membrane (Thermo Fisher). Immunoblotting was performed by using the corresponding antibodies. Horse Radish Peroxidase signals were revealed by using Amersham ECL Prime (Sigma Aldrich). Images were acquired using Chemidoc Imaging system (Bio-Rad). Each band intensity on the immunoblot were quantified using ImagJ software. Quantification was established by the following formula for each viral protein: VLP/(VLP+cell lysate) relative to GAPDH as a loading control.

### Atomic force microscopy on VLPs

Glass-bottom FluoroDish Cell Culture dishes (WPI) were coated with 0.01% poly-L-lysine (Sigma) for 30 min at room temperature and rinsed three times with 1 mL of PBS (ThermoFisher Scientific). Purified MNES, MNE, M(GFP)NES and M(GFP)NE VLPs were diluted 10 times in 100μL TNE buffer and were deposited for 30min at room temperature on the coated dishes for adsorption. Dishes were next filled with 2 mL PBS 1X. AFM imaging was performed at room temperature on a NanoWizard IV atomic force microscope (JPK BioAFM, Bruker Nano GmbH, Berlin, Germany) mounted on an inverted microscope (Nikon Ti-U, Nikon Instruments Europe B.V, Amsterdam, the Netherlands) equipped with a standard monochrome CCD camera (ProgRes MFCool, Jenoptik, Jena, Germany). Fluorescence imaging was performed by wide-field illumination (Intensilight Hg lamp, Nikon) with a 100X objective (Nikon CFI Apo VC, 1.4 NA, oil immersion) and the appropriate filter cube for GFP fluorescence (Ex 466/40, DM 495, BA 525/50). A software module (DirectOverlay, JPK BioAFM, Bruker Nano GmbH, Berlin, Germany) was used to calibrate the tip position with the optical image. AFM topographic images were obtained in TNE using the quantitative imaging (QI) mode with BL-AC40TS cantilevers (mean cantilever spring constant kcant = 0.1 N/m, Olympus). Before each set of acquisitions, the sensitivity and spring constant of the cantilever were calibrated (thermal noise method). The applied force was kept at 200 pN with a constant approach/retract speed of 25 μm/s (z-range of 100 nm). Using the JPK SPM-data processing software, images were flattened with a polynomial/histogram line fit. Low-pass Gaussian and/or median filtering was applied to remove minor noise from the images. The Z-color scale in all images is given as relative after processing. Particle height analysis, based on the height (measured) channel of the QI mode, was performed using the cross-section tool of the analysis software to calculate the maximal central height on each particle.

### Fluorescence Correlation Spectroscopy on fluorescent VLPs

Glass coverslips were coated with 0.01% poly-L-lysine (Sigma) for 30 min at room temperature and rinsed three times with 1 mL of PBS (ThermoFisher Scientific). Purified M(GFP)NES and M(GFP)NE VLPs were diluted 10 times in 100μL TNE buffer and were deposited for 30min at room temperature on the coated dishes for adsorption. Experiments were conducted on a Zeiss LSM780 confocal microscope. The 488nm excitation laser beam is focused onto the sample using a water immersion objective (40x, NA=1.20) resulting in a 210 nm laser waist (ω_0_) in the object plane as calibrated using Rhodamine 6G in water at 25°C (with D=280μm^2^/s). When considering a Gaussian profile for illumination (ω_z_/ω_0_=5), an effective volume (Vo) of ~0.3 fL is obtained, since the sample fluorescence is collected through a confocal pinhole (fixed at 1 Airy unit). Fluorescence intensity fluctuations traces (at least 120) were recorded for 10s and correlated immediately with the built-in Zen software.

Correlograms are then fitted with the PyCorrFit software [20] using a classical 3D free diffusion model

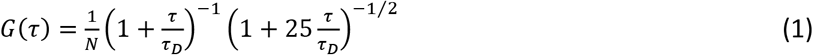

in order to obtain the diffusion time τ_D_ and the number of particle N (eq. 1). All particles were assumed to have similar brightness.

The VLPs concentration (in particle number/ml) is immediately obtained by equation 2

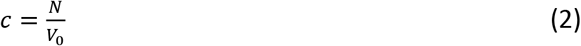

The VLPs diameters (d) were calculated using the following expression:

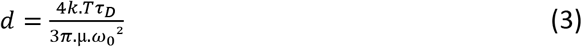

with τ_D_ diffusion time, obtained by fitting the experimental correlograms with eq (1), ω_0_ the laser waist, k the Boltzmann constant, T the temperature in Kelvin and μ the dynamic viscosity of the buffer.

### Photo-Activated Localization Microscopy (PALM) on photoswitchable VLPs

Glass coverslips were coated with 0.01% poly-L-lysine (Sigma) for 30 min at room temperature and rinsed three times with 1 mL of PBS (ThermoFisher Scientific). Purified MN(mEOS2)E VLPs and N(mEOS2) particles were diluted 10 times in 100μL TNE buffer and were deposited for 30min at room temperature on the coated dishes for adsorption. Images were acquired in TIRF mode on an inverted microscope (Nikon Instruments Europe B.V, Amsterdam, the Netherlands) using a NA 1.49 SR-100x TIRF objective at room temperature. A 561 nm laser was used to excite the 405nm photo-convertible mEOS-2 protein. The fluorescence is collected through a dedicated cube containing a dichroic mirror and an emission bandpass filter centered at 617+/-73 nm. 20 000 images were captured by an iXon U897 camera from Andor at a rate of 50 Hz. PALM acquisitions were analyzed using the ThunderSTORM plugin in Fiji [23]. The module DBSCAN of the super-resolution quantification software SR Tesseler [24] was used to quantify cluster diameters.

### Immuno-spotting assay on VLPs

Fluorodishes cell culture (WPI) were coated overnight at room temperature with 0.01% poly-L-lysine (Sigma) and rinsed three times with PBS. Purified fluorescent SARS-CoV-2 VLPs were diluted 10 times in TNE buffer and 100μL were deposited 30 minutes at room temperature on the poly-L-lysine coated dish for adsorption. PFA 4% was deposited 15 minutes at room temperature for fixation. NH_4_Cl was added for 5 minutes at room temperature. Dishes were rinsed once with BSA 3%PBS and were blocked with BSA 3%PBS for 15 minutes at room temperature. Immuno-spotting was performed by using corresponding antibodies. After incubation, dishes were rinsed three times with PBS and stored at 4°C. Then, images of fluorescent VLP were acquired using the TIRF-Microscopy (Nikon) with the 100x objective. 13 Images corresponding to 250-300 fluorescent VLPs were analyzed by ImageJ/Fiji software and ComDet Plugin was used for quantifiying the number of fluorescent VLPs and the double GFP/mCherry colocalization. The image background was estimated at 10% of the fluorescent intensity signal of VLPs and was removed from images before colocalization quantification.

### Fluorescent VLPs Internalization assay

VLPs internalization assays were performed using M(GFP)NE and M(GFP)NES particles. Human pulmonary A549-hACE2 cells were harvested live in cold PBS 1x (after 24h in culture on glass coverslip). M(GFP)NE or M(GFP)NES were diluted (1:20) in transparent DMEM Solution (Gibco) and put into contact with the live cells. The cells were then incubated either at 37°C for 15 min or at 4°C for 1 hour. Unbound VLPs were removed by washing cells 3 times with PBS 1x. The cells were then fixed with 4% PFA and mounted with Prolong™ Gold antifade reagent with DAPI (Invitrogen) with DAPI for confocal imaging. Confocal fluorescence images were generated using a CD7 confocal laser-scanning microscope (Zeiss) equipped with a 50 × 0.5X, 1.2 NA oil objective. For the visualization of images, we have deconvoluted with theoretical PSF. Images were processed with IMARIS viewer, ImageJ/Fiji and Icy for cell center/VLP distance measurements. The cells were imaged as Z stack with 0.3 μm sections.

### Statistical analyses

Statistical analyses were performed using Origin 8.5 and Graph Pad prism software. ANOVA and t-tests were used for statistical comparison of data.

### GraphPad

GraphPad was used for performing graphs.

## RESULTS

### 1- Optimized Virus-Like-Particles production by mimicking SARS-CoV-2 RNA ratio

We first explored the minimal system requirements for SARS-CoV-2 VLPs production. We played with combination of plasmids expressing M, N, E or S (under the same CMV promoter), transfected individually, by pairs or with a plasmid ratio of 3:3:3:5. Transfections were performed on the highly productive HEK293T cell lines, widely used in VLPs production [5] [6] [3] [10] [7]. At 48h post-transfection, VLPs released in cell supernatants were purified by ultracentrifugation over a 25% sucrose cushion in TNE buffer (Figure 1A). Viral proteins in cell lysates and in the purified VLPs were next analyzed by western blot, and the percentage of protein released was calculated (Figure 1B). The results show that only the combination of M, N and E proteins was able to efficiently release VLPs 48h post-transfection. None of the single or dual combinations generated particle production. Without E, some MNS particles were pelletable but the efficiency dropped by 2.5 fold. Without M or N, no VLP were produced in our conditions (Figure 1B). Thus, it appears that the interaction between M and N is absolutely required for VLP production. The combination M, N, E + S could also produce VLPs incorporating S (Figure 1B, central panel), but with a reduction of N proteins in comparison with the MNE combination. In all conditions, cell viability was not impacted (Figure Supplemental 2), indicating that the released particles are not coming from dead cells, and are indeed secreted VLPs. The VLPs production occurred only upon the combination MNE, with or without S, indicating that S is dispensable for SARS-CoV-2 VLPs production (Figure 1B, right panel). Contrary to other recent reported studies [5] [7] [8], ME or MES combinations were not sufficient to form VLPs in our condition. We next evaluated VLPs production using plasmid ratios mimicking the wild-type SARS-CoV-2 mRNA ratio (extracted from RNAseq data of infected cells [17] [18]) coding for the 4 main structural proteins. For that, the expression of M, N, E and S is under the same CMV promotor, with a respective 3:12:2:5 ratio. This transfected plasmid ratio was compared with the arbitrary MNES ratio 3:3:3:5 (Figure 1C). Interestingly, the ratio MNE 3:12:2 was found optimal for the production of SARS-CoV-2 VLPs, with a 1.5-fold increase of M incorporation (Figure 1C, in the graph compare the % of M release 50±4 - lane 2 - with 72±9 – lane 4), but no change in N incorporation (compare the % of N release 54±30 with 55±9), as compared to the MNE 3:3:3 ratio (Figure 1C, graph). The expression of S decreased the amount of N and M proteins incorporated in VLPs by 1.5 fold (Figure 1C, graph, compare the % of M release 72±9 versus 54±15 with S, and, for N, compare 55±9 versus 33±12 with S), suggesting that S modulates the incorporation of N or M most probably due to itself incorporation. Altogether, these data establish that the minimal system for SARS-CoV-2 VLPs production is composed of M, N and E structural viral proteins. The presence of S for VLPs production is optional, but when expressed, S may be incorporated in the VLP accompanied with a diminution of N and M proteins incorporation. However, the incorporation of S on the pelletable VLPs is greater when the transfected plasmid ratio was 3:12:2:5 rather than 3:3:3:5 (see graph Figure 1C, compare the % of S release, 55±9 - lane 3 – to 45±10 – lane 1, respectively). In addition, the maturation of S into S2 was much more efficient when the ratio was respecting the 3:12:2:5 proportionality (Western Blot Figure 1C, compare S2 in lanes 1 and 3 “VLP”). This suggests that the production of mature SARS-CoV-2 particles is most probably fine-tune regulated by viral RNA ratios during infection [17].

The purified MNE and MNES VLPs were next imaged by AFM in buffer for size and shape characterization (Figure 1D), in the conditions previously used for the wild-type virus [19]. MNE and MNES VLPs appeared as spherical particles, with shapes similar to those of WT-SARS-CoV-2 particles. As seen previously reported with the wild-type virus, AFM in liquid in standard scanning mode did not resolved the S proteins at the surface of the MNES VLPs, which has been attributed to the fast dynamics of S in the envelope. The mean height of the VLP was 112±48nm for MNE and 126±17nm for MNES, showing a ~ 25% increase in size as compared with wild-type SARS-CoV-2 particles analyzed by AFM [19] [25] and electron microscopy (EM) [26] [27]. Recent literature on SARS-CoV-2 VLPs reported a size around 100-200nm in diameter either using AFM [7] or EM [5] [6] [7] [9].

### 2- Production of fluorescent SARS-CoV-2 VLPs by tagging M and N proteins without impairing VLP assembly

We next aimed at producing fluorescent SARS-CoV-2 VLPs in HEK293T cells, by co-transfecting a M(Cter-GFP) fusion protein together with M, N and E, supplemented or not with S. M(GFP) incorporation into VLPs collected 48h post transfection was verified by Western Blot in both MNE and MNES transfected cells (Figure 2A). The purified VLPs fraction was deposited on glass coverslips and fluorescent particles were successfully detected using Total Internal Reflexion Fluorescence (TIRF) Microscopy (Figure 2B), for either M(GFP)NE or M(GFP)NES VLPs transfected cells. Fluorescent spots were not detected on the control (Mock) only transfected with M(GFP). Furthermore, correlative fluorescence/AFM imaging confirmed that these fluorescent dots (Figure 2C) are indeed particles of shape and size similar to the unlabeled VLPs (Figure 1D); therefore demonstrating the production of M(GFP)NE or M(GFP)NES VLPs incorporating M(GFP). It is worth noticing that not all the particle found by AFM were GFP(+) indicating that a fraction of the VLPs produced are not labelled.

**Figure 2:**
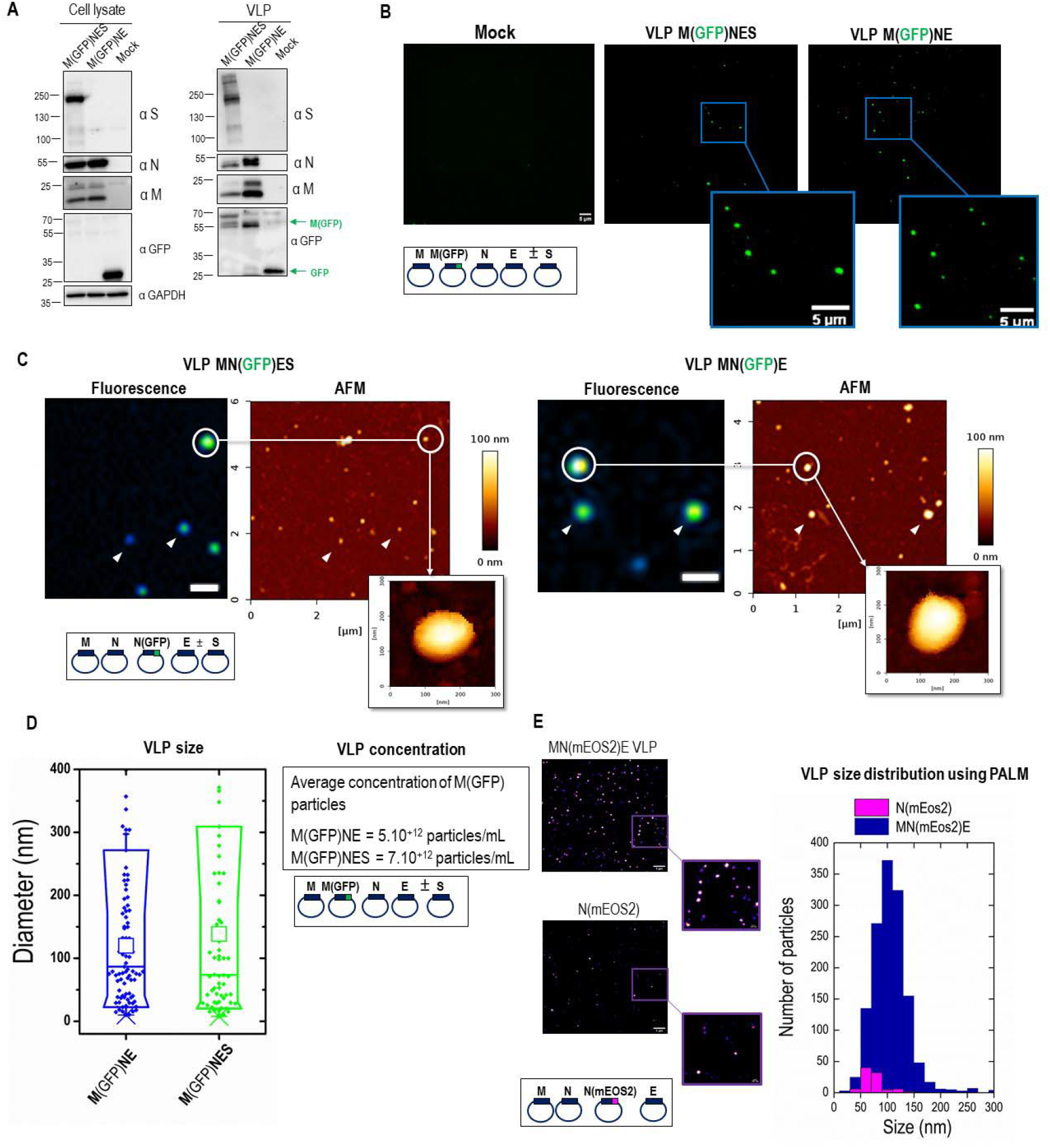
Size and concentration of fluorescent and photoconvertible SARS-CoV-2 VLPs. (A) Western blot of cell lysates and VLPs of HEK293T cells transfected with M/M(GFP), N, E ±S with the optimal plasmid ratio of M/M(GFP):N:E±S as 2/2:12:2±5. (B) TIRF microscopy images of SARS-CoV-2 M(GFP)NE and M(GFP)NES VLPs deposited on a glass coverslip. Scale bar is 5μm. (C) Correlative AFM and wide-field fluorescence images of fluorescent SARS-CoV-2 MN(GFP)E and MN(GFP)ES VLPs. Scale bars represents 1μm. (D) Size and concentration of M(GFP)NE and M(GFP)NES VLPs determined by FCS. (E) Images and quantitative size distribution of MN(mEOS2)E VLPs and N(mEOS2) particles using super-resolution PALM microscopy.

We then determined the M(GFP)NES and M(GFP)NE VLPs size and approximate concentration using Fluorescence Correlative Microscopy (FCS) (Figure 2D). GFP(+) particle concentration of 5.10^12^ to 7.10^12^ particles/ml were respectively found for M(GFP)NES and M(GFP)NE VLPs suggesting a very efficient production considering that not all the VLPs are labelled with GFP. Interestingly, we were able to differentiate VLPs sizes, measuring diameters of 120±10 nm (mean±sd) for VLPs without S and 140±20 nm for VLP with S (Figure 2D). This result is nicely consistent with the average diameter of VLPs obtained by AFM.

In order to accurately measure particle size with single molecule localization microscopy, we next designed and produced mEOS2 photoconvertible fluorescent protein SARS-CoV-2 VLPs using a fusion N(Nter-mEOS2) protein. These photoconvertible VLPs were used to perform Photo-Activated Localization Microscopy (PALM) to provide another estimation of their size [12] [15]. We directly compared these sizes with the particles released into the cell supernatant from cells only transfected with N(mEOS2). The results are shown in Figure 2E. MN(mEOS2)E VLPs exhibited a 108±1 nm (mean±sem) diameter with a high number of particles observed in the field of view (N~1300) while the N(mEOS2) particles were much less numerous (N~60) with a significantly smaller diameter (78±3 nm). This result confirms that SARS-CoV-2 VLPs are in average 100nm in size no matter the technic used thus slightly bigger than the wild-type virus. Secondly, the fact that N alone is being release much less efficiently (20 fold) than the MNE combination confirms that MNE is efficiently producing VLP (Figure 2E, graph of particle size distribution). Overall, our results show the successful incorporation of fluorescent M or N fusion proteins in MNE and MNES VLPs, and show that this incorporation does not impair VLPs assembly. Dual color fluorescent labeled VLPs were next tried and produced by co-expression in HEK293T cells of M(Cter-mCherry) and N(Nter-GFP) with M, N and E supplemented or not with S. M(mCherry) and N(GFP) incorporation into VLPs collected 48h post transfection was verified by Western Blot (Figure 3A). Figure 3A exhibits 2 bands at molecular weights of approximately 80 and 105 kDa, corresponding to respectively fluorescently (GFP or mCherry, ~26kDa) tagged M (~55kDa) and tagged N proteins (~80kDa). Two color TIRF images of these dual color VLPs showed the presence of a mix of GFP(+) or mCherry(+) single labelled VLPs as well as particles containing the two fluorescent labels (Figure 3B). A total of 250-300 fluorescent particles were counted over 13 images (frames) on which we evaluated the percentage of particles only mCherry(+) or only GFP(+). The results showed that half of the fluorescent VLPs were N(GFP)(+) and half were M(mCherry)(+) in the case of MNE VLPs but only one third were N-GFP(+) in the case of MNES VLPs, indicating that upon S incorporation in the VLPs, less N(GFP) was incorporated. The colocalization of the dual color GFP/mCherry VLPs was also quantified, which allowed us to monitor the relative incorporation of M(mCherry) and N(GFP) in a VLP (Figure 3C). We quantify that only 20±3% and 11±7% of the VLPs are dual color containing M(mCherry) and N(GFP) in the same MNE and MNES VLPs respectively (Fig3C) when mixing label and unlabeled proteins. This indicates that VLPs could incorporate the two fluorescent labelled M and N proteins without impairing assembly. We also tried to produce M(mCherry)N(GFP)E VLP, i.e. in the presence of both tagged M and N, but this was totally preventing VLPs production (data not shown). Since the fluorescent protein tag is located at the C-ter of N and M, it should be located inside the VLP in the case of N and outside in the case of M. Only when mixing both label and unlabeled proteins at a ratio of 1:1 for M or at a ratio of 3:9 for N(GFP) versus N, then production of labelled VLPs was possible, compatible with VLPs assembly.

**Figure 3:**
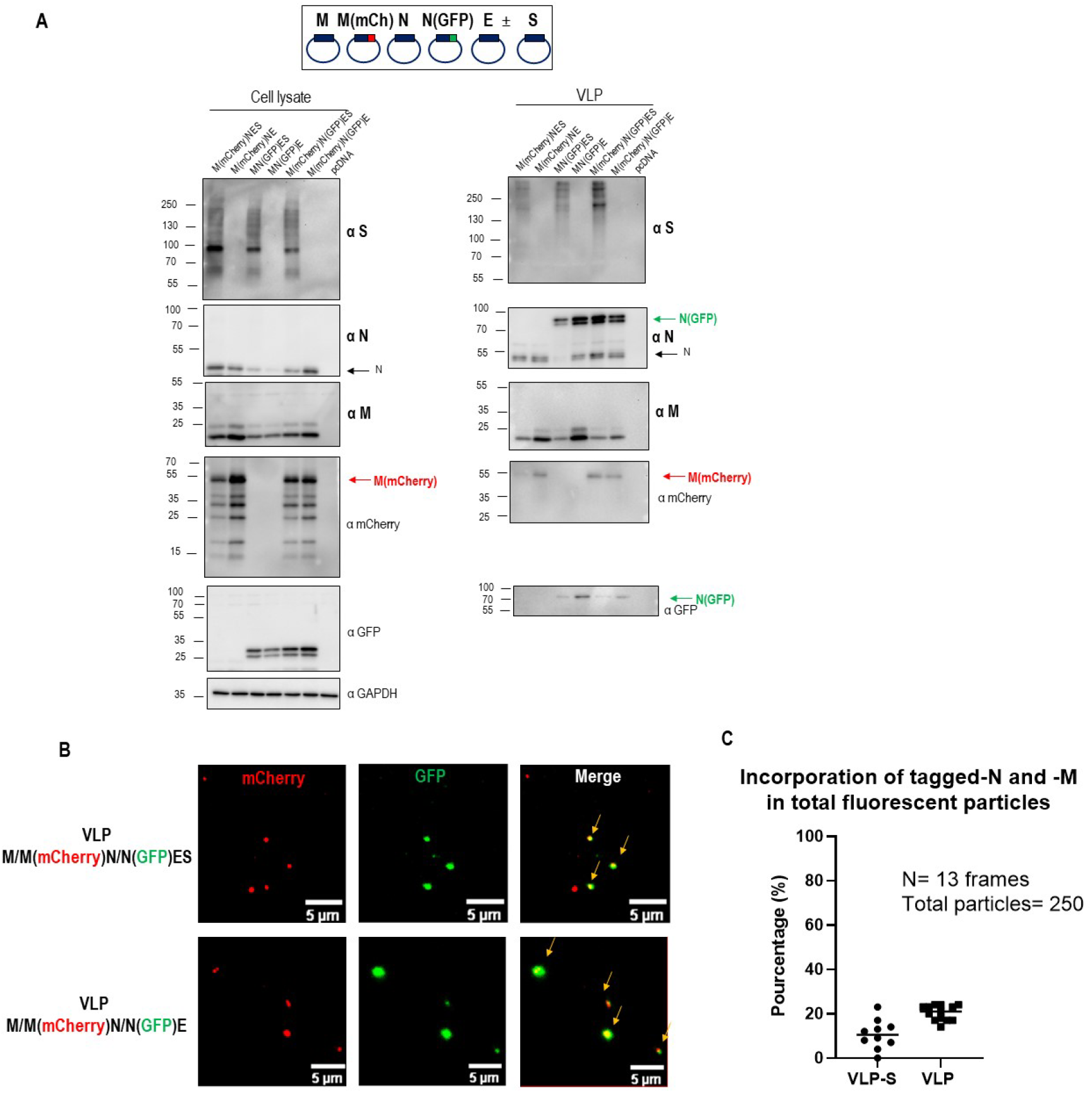
Description of dual-color fluorescent SARS-CoV-2 VLPs with tagged-M and -N incorporation. (A) Western blot of cell lysates and VLPs of HEK293T cells transfected with M, M(mCherry), N, N(GFP), E and S with the optimal ratio of M/M(mCherry)/N/N(GFP)/E±S plasmids 2/2:9/3:2±5. (B) Images of dual color SARS-CoV-2 M(mCherry)N(GFP)E±S VLPs using TIRF Microscopy. Arrows show dual color VLPs. Scale bar is 5μm. (**C**) Quantitative colocalization of dual color GFP/mCherry VLPs using ComDet plugin in ImageJ. For each conditions, 250-300 fluorescent particles were counted over 13 images (frames). One point in the graph represents the % of GFP/mCherry colocalization in one frame.

### 3- Imaging S incorporation in fluorescent VLPs

The M(GFP)NES fluorescent VLPs allowed us to quantify the percentage of GFP labelled VLP incorporating S by immuno-spotting coupled to TIRF microscopy, using an anti-S neutralizing antibody and an AlexaFluor555 secondary antibody (Figure 4A). Non-specific binding of the anti-S antibody/secondary antibody was 6±6% on the M(GFP)NE VLPs, while absence of VLP did not show any significative background (less than 1% for “no VLP”). Quantification of fluorescent red/green dot spots colocalization indicated that 27±12% of the M(GFP)NES VLPs incorporated S (Figure 4B). Although, since some VLPs were not GFP labelled, “red” dots also appeared. In order to prove that these dots were more likely VLPs rather than exosomes/extracellular vesicles (EVs) containing S, we checked the colocalization between these “red” S dots and CD81 with an anti-CD81 exosomal marker (Figure Supplemental 3) using the same immuno-spotting technique. Less than 10% of the “red” dots containing only S were colocalizing with CD81 and none of the M(GFP)NES VLPs were CD81 positive, suggesting that M(GFP)NES VLPs are a mixture of GFP(+)VLPs and unlabeled VLPs containing S but not CD81 EVs. Our results show that the Spike S is well incorporated on MNES VLPs with or without M-GFP (Fig4B), suggesting that S incorporation is mainly driven by MNE confirming the results seen by immunoblotting (Figure 1B). We also checked that these VLPs were distinct from CD81(+) extracellular vesicles (Figure Supplemental 3). On the other hand, not all the VLPs-(GFP) were carrying the Spike, suggesting that part of SARS-CoV-2 particles assemble without S.

**Figure 4:**
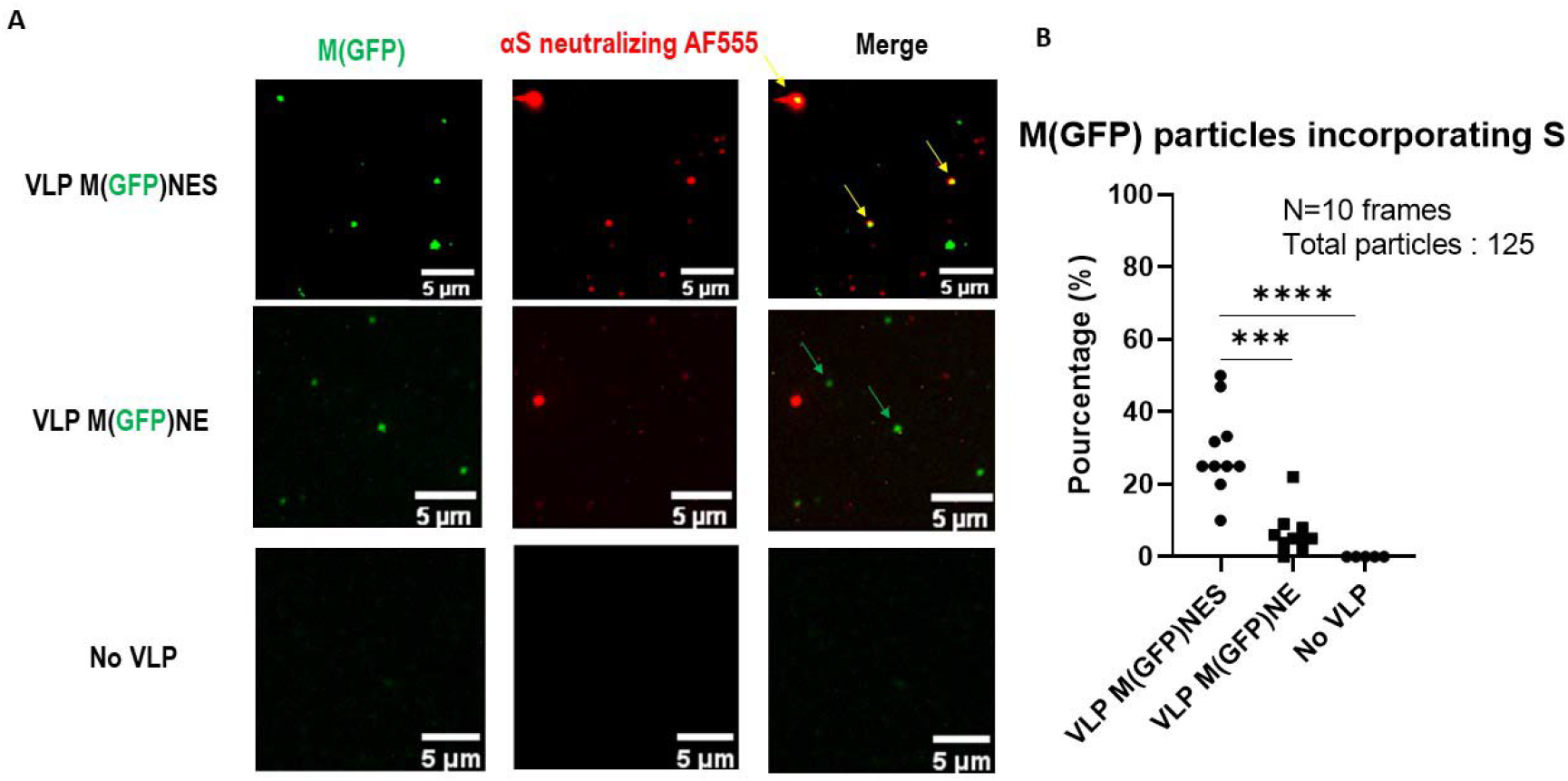
Incorporation of the Spike S on fluorescent SARS-CoV-2 M(GFP)NES VLPs using immuno-spotting coupled to TIRF-Microscopy. (A) TIRF images of M(GFP)NES VLPs, M(GFP)NE VLPs and control. An anti-S antibody coupled to a AF555 secondary antibody were used for S immunolabeling. Yellow arrows show the anti-S antibody-AF555 bound tp VLPs-(GFP). Green arrows show VLPs-(GFP). Red arrows show antibody-labelled VLPs. Scale bar is 5μm. (B) Percentage of incorporation of S on M(GFP)NES, M(GFP)NE and control. For each condition, 125 fluorescent particles were counted over 10 images (frames). One point in the graph represents the % of S incorporation in the VLP(GFP), i.e. % of colocalization in each frame. t test was calculated and p-value between VLP M(GFP)NES and VLP M(GFP)NE is 0.0001 and p-value between VLP M(GFP)NES and No VLP is <0.0001.

### 4- VLP-Spike internalization in host pulmonary cells expressing ACE2 receptor

In order to study the functionality of the Spike incorporated in the VLPs, we performed an internalization assay of the fluorescent SARS-CoV-2 VLP-(GFP). To do so, fluorescent M(GFP)NE VLPs versus M(GFP)NES VLPs were incubated with pulmonary A549 cells expressing the human receptor hACE2. The internalization of both fluorescent VLPs, M(GFP)NE versus M(GFP)NES, was monitor at 37°C (active endocytosis) and at 4°C (blocked endocytosis). A difference of internalization between M(GFP)NES VLPs) and M(GFP)NE VLPs was revealed using quantitative confocal microscopy (Figure 5A). To quantify this difference, we measured the distances (d_1_,d_2_,…d_n_) of VLPs, identified as GFP bright dots, to the center of the cell (Figure 5B), this cell center being the intercept of the major and minor axis (r_1_ and r_2_), established for each cell. As these radii represent the mean longest (r_1_) and shorter (r_2_) distances between the cell border and the cell center, it suggests that every VLP distance that will be found in between these two radii corresponds to a VLP located at the border of the cell. On the opposite, every cell center-VLP distance found below r_2_ indicates that the VLP is inside the cell. We then plot the distribution of the distances (n>100) for each different condition and plot the average value of the major axis, r_1_=17.6+/-3.7 μm (mean+/− sd) and the minor axis, r_2_=11.6+/-3.6 μm (mean+/− sd) of the different cells (n=16) (Figure 5C). In Figure 5C, it is immediately seen that in absence of S, the average M(GFP)NE VLP-cell center distance is found in between r_1_ and r_2_, both at 4°C (d=13.4+/-3.2 μm, mean+/− sd) and at 37°C (d=13.5+/− 3.5 μm), these mean values being not statistically different (p=0.98, Mann Withney U test). For MNES VLP-M(GFP), at 4°C, we observed a slight but statistically significant (0.01<p<0.04) decrease in the mean VLP-cell center distance (d=11.8+/-5 μm) when compared to M(GFP)NE VLP. From figure 5C it can also be seen that, in this case, the distance distribution is widened and that some VLPs start to exhibit distances at value much below r2, suggesting entry of these VLPs. This is confirmed when observing the M(GFP)NES VLP-cell center distance distribution at 37°C. In this case the mean distance value is approximately divided by two (d=6.1+/-3.0 μm) and is statistically strongly different form all the other distributions (10^-33^<p<2.10^-15^) we established here. This clearly suggest that the presence of S on the surface of the VLPs induce their endocytosis in A549-hACE2. To confirm that this endocytosis was mediated by hACE2, we expressed in A549 cells a mScarlet1 tagged version of ACE2 and repeated the same experiment of M(GFP)NES VLPs internalization for 15min at 37°C on live cells. After fixation, z-projection imaging of M(GFP)NES VLPs in A549-hACE2mScarlet1 cells showed the internalization of the M(GFP)NES VLPs on the receptor ACE2. Z-projection imaging at 4°C showed reduced internalization of the M(GFP)NES VLPs decorating the cell surface on the hACE2mScarlet1 receptor (Figure 5D). The mScarlet1 versus GFP fluorescence emission is shown as a control (Figure Supplemental 4). Quantification of confocal images reveal that more than 65% of the GFP(+) dots were associated with the mScarlet1(+) dots (Figure 5D), reinforcing the idea that all the VLPs containing S were able to load on the ACE2 receptor at the cell surface of pulmonary cells mimicking hACE2-dependent SARS-CoV-2 endocytic entry. A colocalization of ~80% of the M(GFP)NES VLPs on the receptor hACE2mScarlet1 was observed (Figure 5E). In the absence of VLP, the cellular distribution of hACE2mScarlet1 remains diffuse (Figure 5F).

**Figure 5:**
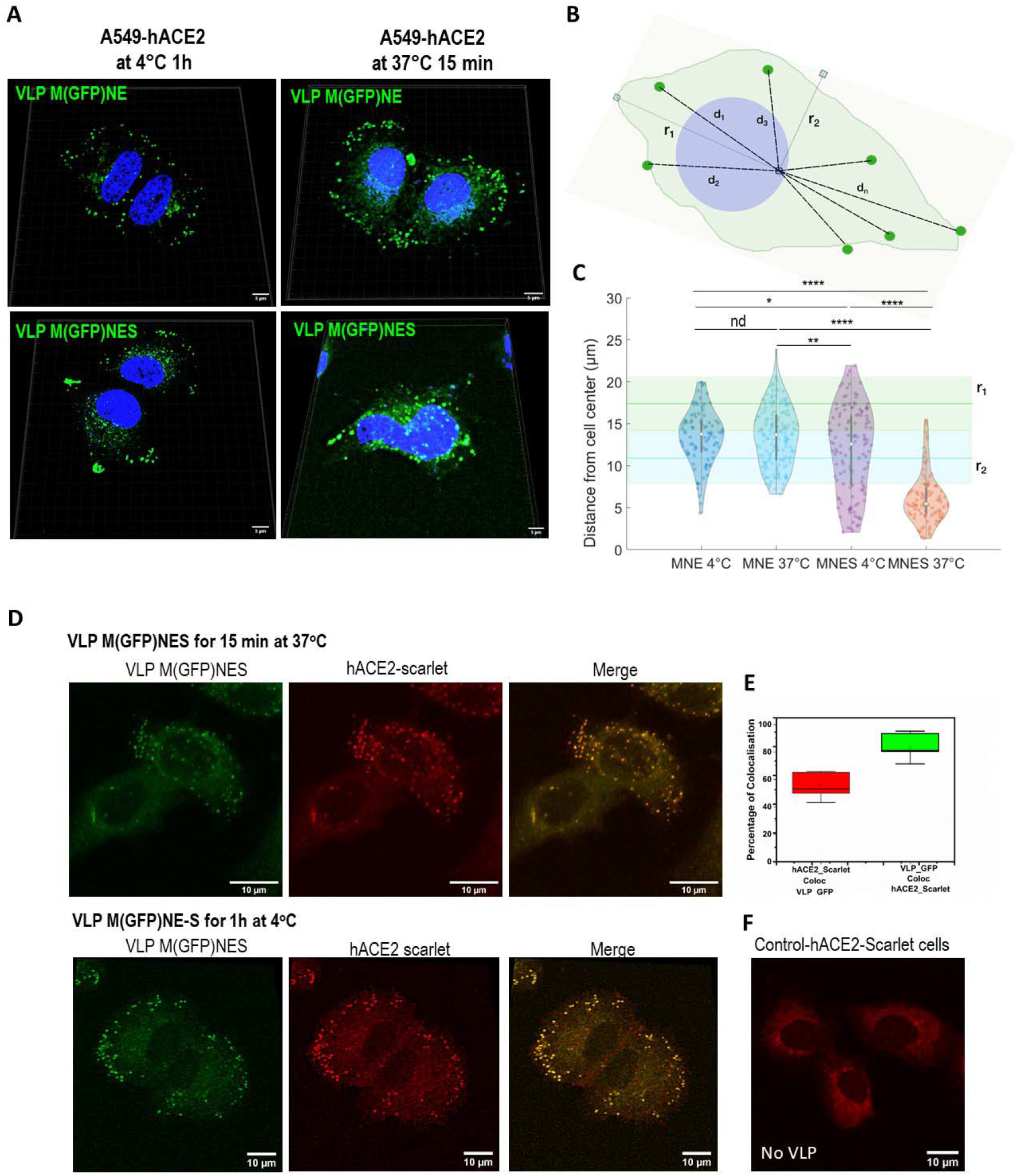
Internalization of VLPs-(GFP) with and without Spike on A549 pulmonary cells expressing hACE2 using confocal fluorescence microscopy. (A) M(GFP)NE±S VLPs were incubated on A549-hACE2 cells at 37 °C for 15min or at 4°C for 1h. Nuclei were stained with DAPI. (B) Schematic of distance measurement and cell center determination from minor and major axis of the cell. (C) Violin plot of the distribution of the VLP-cell center distances for the different conditions (MNE vs MNES at 4°C and 37°C respectively). The average value and the standard deviation of the major and minor axis (r_1_ and r_2_) observed in the different cell conditions (n=16 cells) are respectively plotted as green and blue lines (mean) surrounded by rectangle (sd). All the conditions except MNES at 37°C have distances mean values lying between these two limit axis values. * p-value < 0.05; ** p < 0.01; **** p < 1.e10^-15^; nd non significative. (D) Confocal images of M(GFP)NES VLPs (in green) and A549-hACE2-mScarlet1 (in red) at 37°C (upper panel) and at 4°C (lower panels) showing internalization of VLPs at 37°C (E) Percentage of colocalization of hACE2-mScarlet1 in M(GFP)NES VLPs (in red) and of M(GFP)NES VLPs in hACE2-mScarlet1 (in green) showing that most of the labelled VLPs are surrounded by hACE2-mScarlet1 receptor. (F) Image of a cell expressing hACE2-mScarlet1 without VLPs (control)

## DISCUSSION

As the SARS-CoV-2 pandemic persists around the world, it is necessary to further gather knowledge on this virus, especially on developing tools for studying virus assembly and virus entry in a safe BSL-2 environment (outside a BSL-3 and with non-infectious particles). Therefore, Virus-Like Particles production and characterization are essential to fully apprehend SARS-CoV-2 assembly mechanisms, virus entry or to be used for the development of VLP-based vaccines. In this study, we established a minimal system for SARS-CoV-2 based VLPs production, requiring MNE/S with an optimized transfected plasmid ratio based on the RNAseq ratio measured in infected cells [17] [18]. We show that the transfection and expression of M, N, E, or S alone are not sufficient to produce VLPs, as well as the combination of only two structural proteins. Our conclusion is that M, E, and N is the best condition, at a 3:12:2 ratio, required for optimal VLPs production (as shown by Kumar et al. [10] and Boson et al. [14]). We also observed that the addition of S reduces the incorporation of N and, to a lesser extent M (as seen by immunoblots, Figure 1 and by dual color labelled VLP, Figure 3) suggesting that the incorporation of the transmembrane Spike in the VLPs reduces N incorporation in favor of S. Moreover, S incorporation was favored in the optimized plasmid ratio and its maturation was much greater. Apparently, the presence of the packageable genomic RNA was not required for VLPs formation since all these VLPs were produced without. Some of these findings could be in discordance with other studies. As a matter of fact, Xu et al. [5] showed that M alone is able to exit the cell, with this protein being the driver and M and E the minimal system. Indeed, in Figure 1B, we can notice some M released from the transfected cells but nothing compare to the MNE combination. Plescia et al. [6] demonstrated that after being transfected alone, M is not present in the cell supernatant, as for M and E combination. However, they showed that the minimal requirement to obtain both proteins in the supernatant is to combine M with N. They also argue that the most efficient production is achieved with an M, N, and E combination, as the addition of E is playing an important role in particles assembly and release (see Bracquemond et al. [4]). In this study, the addition of S is not changing the proportion of M or N, which differs from Xu et al.[5]. Finally, other teams concluded that SARS-CoV-2 VLPs production was sufficient when VLPs are based on M, E, and S combination in mammalian cells (Swann et al. [7]) or in insect cells (Mi et al. [8]). Our study shows that MN is required and that the addition of E makes the yield of VLPs production 2.5 fold better. A combination of M, N, and E was recently also reported for producing SARS-CoV-2 VLPs in plants (Moon et al. [9]) and previously in mammalian cells in the case of SARS-CoV [12].

With the intention of optimizing both VLPs formation and production, we performed an assay with an optimal ratio of the different structural proteins based on the viral RNA ratio coding for the 4 main structural proteins found in infected pulmonary Calu3 or intestine Caco2 cells [17] and in human pulmonary A549hACE2 cells [18]. Consequently, RNAseq data analysis allowed us to establish a plasmid ratio of 3:12:2:5 for M:N:E:S. This ratio increases VLPs production by 1.5 fold for MNE and MNES, as compare with the 3:3:3:5 ratio, with more S incorporation and more mature S2 on the VLP (Figure 1C), mimicking the wild-type virus.

We confirmed that particles analyzed by AFM were genuinely VLPs and rather not extracellular vesicles (Figure 2C). To confirm that aspect, we checked one exosomal marker, CD81, using immunospotting on GFP-VLP and confirm very low (less than 5%) or no colocalization between VLP and CD81(+) EVs (Supplementary Figure 3). This suggests that SARS-CoV-2 VLPs are exiting through a secretory pathway that is different from EVs.

While Swann et al. [7] reported a 200 nm diameter and a height of 50-60 nm for MES VLPs using AFM, we measured a size of 126±17nm for MNES VLP using the same technique (Figure 1D). No significant differences were observed between MNE, MNES and the wild-type virus (Lyonnais et al. [14]) by liquid AFM imaging. To confirm the size of these particles and to allow the study of SARS-CoV-2 using fluorescence and single molecule localization microscopy, we produced VLP incorporating M or N proteins in fusion with fluorescent or photoconvertible proteins. Using FCS and PALM, we found a 120-140nm particle size range (Figure 2D,E): 100-110 nm for MNE VLPs, and 130nm for MNES VLPs, which is consistent with AFM results (Figure 1D). It is to notice that VLPs size are slightly bigger than wild-type virus particles as it has been described using AFM [19], TEM [19] or cryoET [27] [28] [29]. However, Spikes are difficult to observed: MNE and MNES VLP size could only been distinguished using FCS, where 2 populations were observed: one bigger than 100nm in diameter, and one equal to 100nm in diameter, suggesting that SARS-CoV-2 is producing some particles with and without Spikes as seen by immuno-spotting (Figure 4). Using electron microscopy, Swann et al. [7] established a size around 130 nm while Xu et al. [5] obtained sizes between 70-90 nm depending on the cell lines used for VLPs production; Mi et al. reported around 100nm [8] and Moon et al. [9] around 75nm while we observed 90nm for the wild-type virus [14]. These differences between our data and the ones from electron microscopy could arise from the dehydration process required during TEM sample preparation or from the absence of genomic RNA shaping the inside nucleocapsid N-RNA core that might change particle assembly. The incorporation of other viral or cellular proteins could also be considered for tuning particle assembly. In another study, McMahon et al. [15] used high-throughput super-resolution microscopy to measure the size of the SARS-CoV-2 particles, showing that the virus’s diameter distribution is mono-dispersed and centered at 143 nm. This size is more in accordance with the one we found using FCS for the measuring M(GFP)NES VLPs size in solution (Figure 2D).

Labelling of VLPs with M Cter- and N Nter-tagged fluorescent proteins (Figures 2 and 3), appeared possible only if there was a mixture of labelled and unlabeled M and N proteins during particle assembly. This could be very useful tools for visualizing virus assembly or entry in target cells as reported recently [6]. In addition, following double fluorescently labelled VLP could be an advantage in imaging VLP overtime in different cell compartments, for example.

Concerning Spike incorporation in the VLP, thanks to immuno-spotting with a neutralizing S antibody, we concluded that at least 25% of the M(GFP)NES VLPs incorporate the spike. This number might be under-estimated due to the deposit of the MNES VLPs on the glass surface and the affinity of the S antibody to the Spike, but it tends to support the observation that not all the VLPs produced by the mammalian cells are incorporating the spike. It would be interesting to vary the ratio of Spike expression in the productive cells to see if it would change the level of incorporation in the VLPs, and to try different anti-S neutralizing antibodies. Our results also suggest that the injection of MNES VLP mimicking the virus to animal models should provide adequate antibody responses without being infectious.

Overall, we can conclude that the optimized SARS-CoV-2 based VLPs mimics the assembly of SARS-CoV-2 particles.

Finally, our study has shown that in the presence of overexpressed hACE2 receptors at the cell surface, M(GFP)NES VLPs can recognize the receptor and being endocytosed, confirming that the Spikes incorporated in the VLPs are functional in recognizing the receptor ACE2. Interestingly, a recent article [29] is demonstrating that this type of VLPs can package viral RNAs to transduce and express genes in target cells, which reinforces the strong potential of SARS-CoV2 VLPs in vaccine development.

Overall, we report that the minimal system required for VLPs formation is M, N, and E structural proteins incorporating S and that VLPs can be optimized in production and maturation by mimicking SARS-CoV-2 mRNA ratio in infected cells. While VLPs could be used to study the viral life cycle of SARS-CoV-2 or as vaccines prototypes, tagging VLPs with fluorescent proteins could be very useful to image SARS-CoV-2-ACE2 entry and particle assembly using fluorescence nanoscopies.

## ACKNOWLEDGMENT

We are grateful to Dr Olivier Schwartz (Institut Pasteur Paris France) for the gift of the hACE2-mScarlet1 expression plasmid. Confocal and TIRF microscopy were performed at the Montpellier Imaging Center for Microscopy (MRI) and the correlative fluorescence Atomic Force Microscopy at CEMIPAI CNRS Montpellier, France, which was funded by the REDSAIM program (Montpellier University).

## AUTHOR CONTRIBUTIONS

MG, CA, PM, DB, JS, AM, SL performed experiments. PM generated A549hACE2 and A549hACEmScarlet1 cell lines. MG directed VLPs production and purification, immunoblot, immuno-spotting and data analysis; SS generated M, N, E, S M(GFP) and M(Cherry) plasmids; DB generated N(mEOS2) and N(GFP) plasmids; CA and SL performed Bio-AFM and image analysis of VLPs; MG performed TIRF-M VLPs imaging and analysis; AM and CF performed live-VLPs fluorescence FCS and PALM microscopy study and analysis. DM and JS performed VLP-cell entry confocal microscopy study and analyzed data with the help of CF. DM conceived, directed, and supervised the study. MG and DM wrote the manuscript. MG, DM, CA, JS, SL and CF edited the manuscript. SS and DM raised funding.

## SUPPLEMENTARY DATA

**Supplementary Figure 1:**
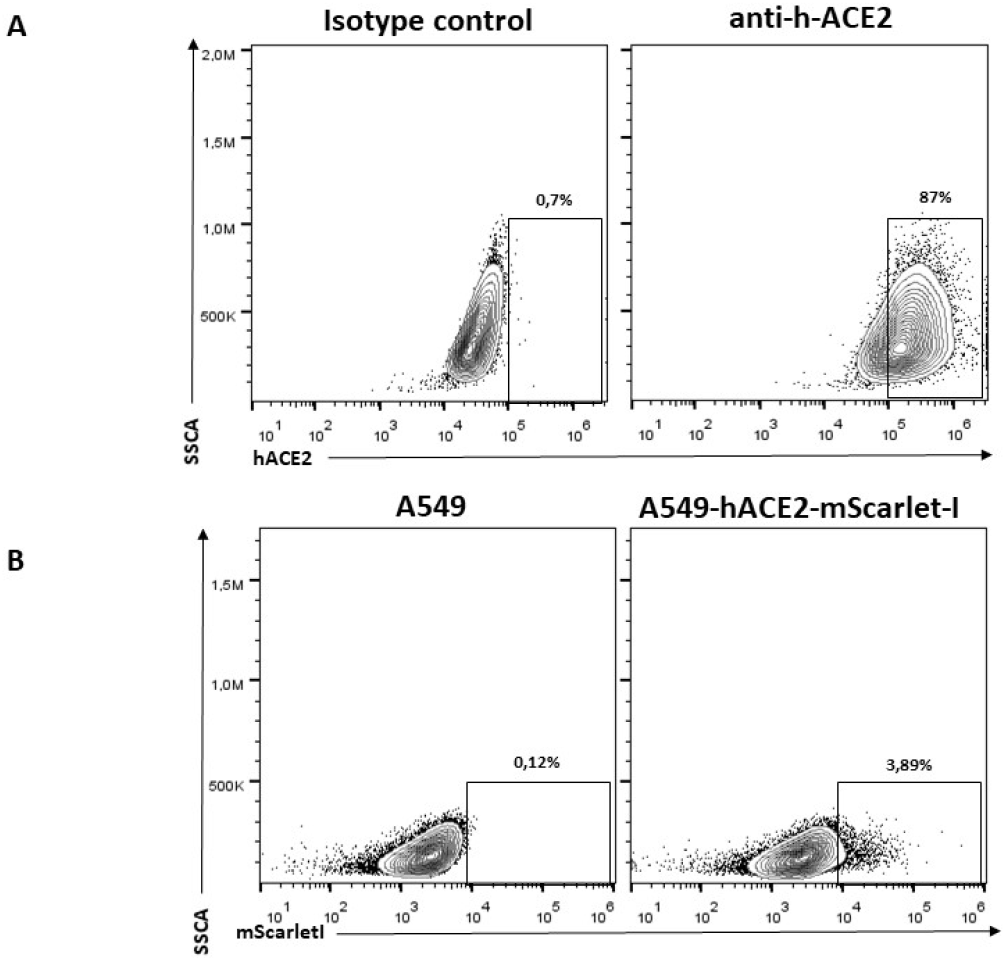
Detection of hACE2 and hACE-mScarlet1 at the cell surface of the A549-hACE2 and A549-hACE2mScarlet1 pulmonary cell lines using flow cytometry. (A) Dot plot representing side scatter (SSC) in function of mean intensity (secondary AF488 antibody). Gate represents the pourcentage of hACE2 positive cells. Cells labeled with only secondary antibody are used as control. (B) Dot plot representing side scatter (SSC) in function of mean intensity of mScarletI (Ext569/Em594nm). Gate represents the pourcentage of mScarletI positive cells. Non transduced A549 cells are used as control.

**Supplementary Figure 2:**
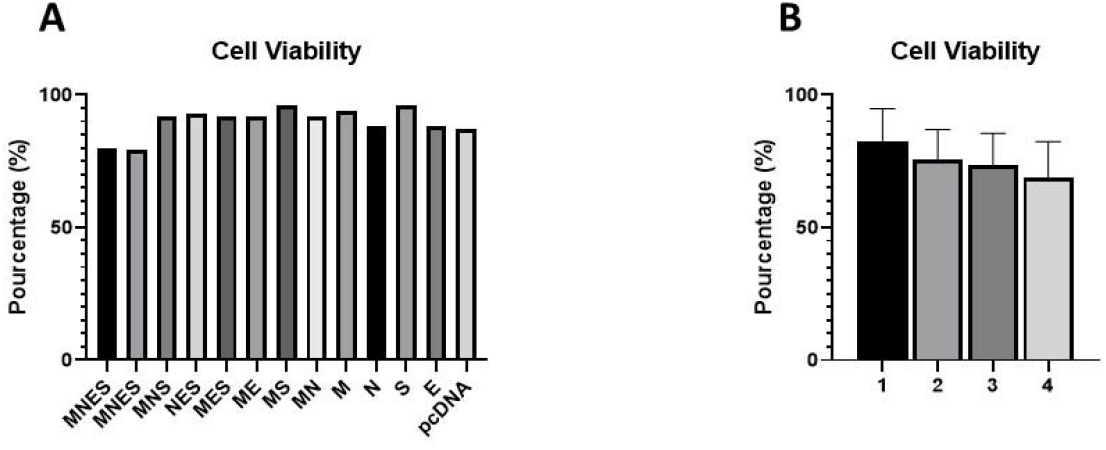
Cell viability of transfected HEK293T cells when producing VLPs for different experimental conditions. (A) Cell viability of assays based on constant ratio corresponding to Figure 1B. (B) Cell viability of assays based on mimicking WT SARS-CoV-2 mRNA ratio corresponding to Figure 1C.

**Supplementary Figure 3:**
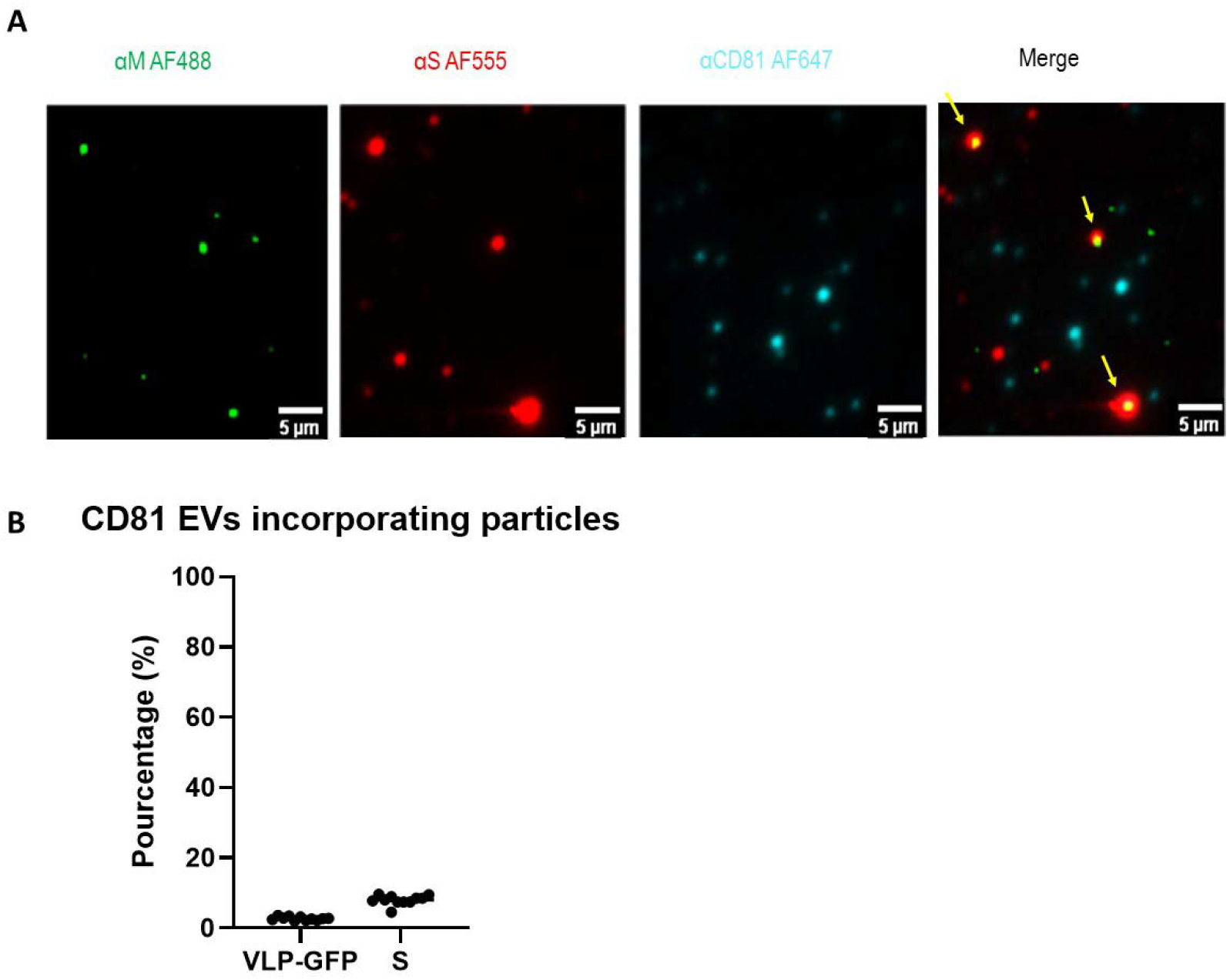
CD81(+) EVs and SARS-CoV-2 M(GFP)NES VLPs are distinct entities as revealed by immuno-spotting coupled to TIRF-M. (A) Imaging of incorporation of the M(GFP)NES VLP on CD81-exosomes labelled with M(GFP), with a neutralizing antibody anti-S coupled with secondary AF555 antibody and an antibody anti-CD81 coupled with secondary AF647using immuno-spotting coupled to TIRF-M, showing that M(GFP)NES can contain CD81 but CD81 exosomes are not containing M(GFP) or S. Scale bar is 5μm. (B) Percentage of incorporation of M(GFP) on CD81(+) EVs showing that CD81(+) exosomes are not containing M(GFP) or S. Scale bar is 5μm.

**Supplementary Figure 4:**
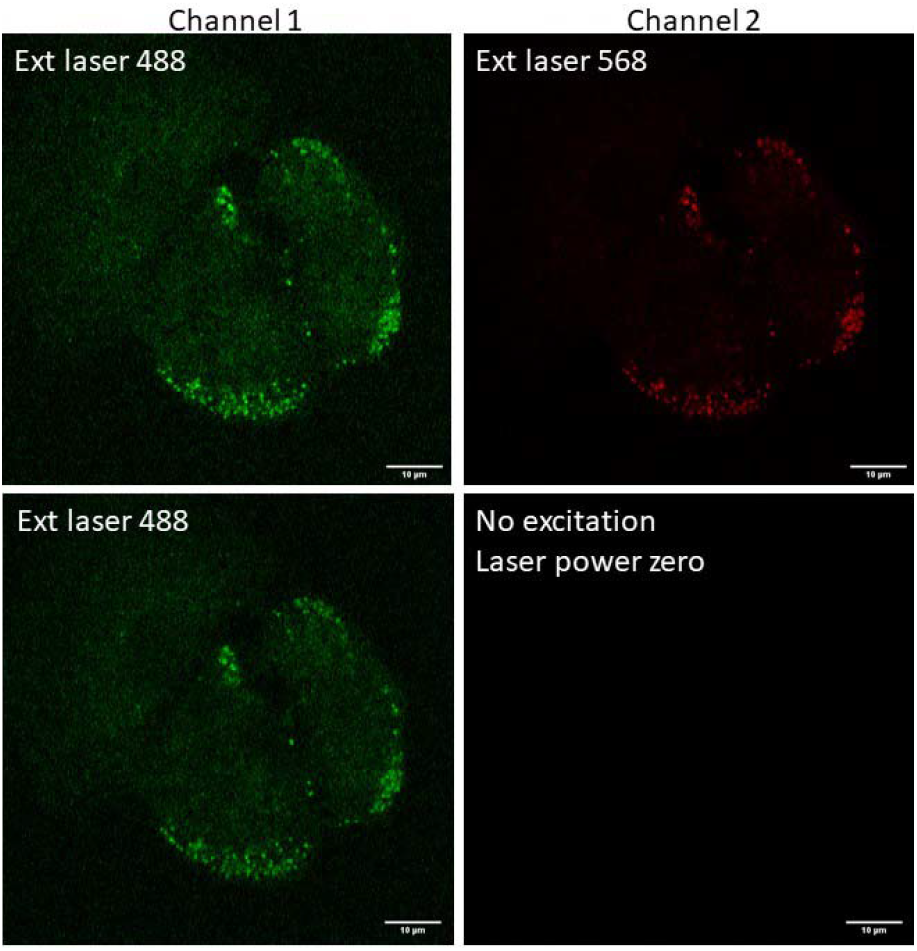
Control imaging of VLP(GFP) in A549-hACE2mScarletl cells using confocal laser microscopy. Confocal images of VLP(GFP) on A549hACE2mScarlet1 cells showing that the GFP fluorescence (488nm excitation) is not emitting into the mScarlet1 channel (561nm excitation). Without excitation of mScarlet1 the “red color” dots do not appear on the image.

